# A domestic cat whole exome sequencing resource for trait discovery

**DOI:** 10.1101/2020.06.01.128405

**Authors:** Alana R. Rodney, Reuben M. Buckley, Robert S. Fulton, Catrina Fronick, Todd Richmond, Christopher R. Helps, Peter Pantke, Dianne J. Trent, Leslie A. Lyons, Wesley C. Warren

## Abstract

Over 94 million domestic cats are considered pets, who, as our companions, are also susceptible to cancers, common and rare diseases. Whole exome sequencing (WES) is a cost-effective strategy to study their putative disease-causing variants. Presented is ~35.8 Mb exome capture design based on the annotated Felis_catus_9.0 genome assembly, covering 201,683 regions of the cat genome. WES was conducted on 41 cats from various breeds with known and unknown diseases and traits, including 10 cats with prior whole genome sequence (WGS) data available, to test WES capture probe performance. A WES and WGS comparison was completed to understand variant discovery capability of each platform. At ~80x exome coverage, the percent of on-target base coverage >20x was 96.4% with an average of 10.4% off-target. For variant discovery, greater than 98% of WGS SNPs were also discovered by WES. Platform specific variants were mainly restricted to a small number of sex chromosome and olfactory receptor genes. Within the 41 cats with ~31 diseases and normal traits, 45 previously known disease or trait causal variants were observed, such as Persian progressive retinal degeneration and hydrocephalus. Novel candidate variants for polycystic kidney disease and atrichia in the Peterbald breed were also identified as well as a new cat patient with a known variant for cystinuria. These results show the discovery potential of deep exome sequencing to validate existing disease gene models and identify novel gene candidate alleles for many common and rare diseases in cats.

## Introduction

Precision / Genomic medicine is the next frontier to conquer in veterinary medicine, however, the appropriate resources are necessary for robust implementation of genomic medicine in clinical practice^1^. One tool, which has been successfully applied to the diagnosis of rare diseases in humans, is whole exome sequence (WES) analysis, a cost-effective method for identifying potentially impactful DNA variants in the coding regions of genes^2^. Alternatively, whole genome sequencing (WGS), captures DNA variants spanning the entire genome for which the vast majority have an unpredictable impact. The increased cost of WGS analysis raises the question whether, WES, the more affordable option, can be just as effective for diagnosing novel disease variants in cats?

Over the last decade, a surge of studies using next generation sequencing, in particular WES, has led to many novel discoveries in disease causation. WES became recognized as a more efficient means for genome resequencing in 2007 and has increasingly been used to help diagnose patients with rare and genetic diseases^3,4^. By selectively sequencing all protein-coding regions to great depth, WES is a dependable method to find exome variants^5^. Most often WES is a powerful approach to study Mendelian inherited diseases because highly functional impact variants rarely appear in the healthy populations^6 7^. In humans, exome sequencing has been used to study a wide-range of diseases, cancers, and analysis of autism spectrum disorder^8,9,10^. The discovery of exome variants has led to therapeutic targets for drug development, and genetic markers for innovative clinical applications in companion animals and humans^11,3^. The exome-only approach is especially successful in cancer studies by cost-effectively providing variant information about the normal and tumor genomes within patients, supporting the identification of tumor drivers that may indicate a druggable potential for therapy^12^.

Exome sequencing has also proven successful in various species. In mice, exome data has been used to study Mendelian inherited disorders, and to complete a cross-species analyses with humans^13,14^. In 2014, the first dog exome capture study demonstrated that provided sufficient coverage, an average of 90% of bases and targets covered, causative allele discovery has great potential^15–18^. Since the development of the dog WES capabilities, several studies have been successful in identification of causal variants for various diseases including a two base pair deletion in *SGCD* for muscular dystrophy, and a splice site variant in *INPP5E* in dogs with cystic renal dysplasia^19,20^. The domestic dog with a large number of isolated breeds are an important genetic resource for cancer studies, and WES has shown similar oncogene variant patterns that enable comparative analysis in humans^21,22^. On the other hand, an analysis of human and canine bladder cancer using WES data identified novel mutations in *FAM133B*, *RAB3GAP2*, and *ANKRD52* that are unique to canine bladder cancer suggesting biological differences in origin^23^.

Cats have long been recognized for their potential in modeling some human diseases, such as retinal blindness and testing for clinical therapies^24,25^. In domestic cats, approximately 150 variants are associated with over 100 genetic traits or diseases, many as biomedical models for human diseases^26^. As feline genomic resources continue to advance, greater numbers of diseases caused by single base variants are being discovered, such as two novel forms of blindness in Persians and Bengal cats^27,28^.

This study outlines the development and performance of cat WES approaches and demonstrates its use to identify putatively causative alleles for disease and normal trait phenotypes. Our WES analyses included 41 cats with phenotypes in which causative alleles are known and/or unknown, leading to the discovery of three novel, likely causal, disease variants and the confirmation of a variety of known diseases and traits and their population allele frequencies. These results prove the discovery potential of feline WES to validate existing disease gene models and identify new ones.

## Methods

### Exome design

The annotated exons from the Felis_catus_9.0 reference genome assembly were used as the basis to design the exome capture probes^29^, incorporating the NCBI RefSeq release 92 annotation. The coding sequences for the primary chromosomes were extracted and consolidated into a non-overlapping set of features, totaling 35,855,889 bases divided over 201,683 regions. Since Y chromosome genes are not represented in the Felis_catus_9.0 reference, a set of coding sequence features from the *Felis catus* Y chromosome genomic sequence (NCBI accession KP081775) was used^30^. The cat exome panel was designed by Roche Sequencing Solutions (Madison, USA)^31^. A capture probe dataset was constructed for the full cat genome by tiling variable length probes, ranging from 50 - 100 bases in length, at a five-base step across all sequences. Each capture probe was evaluated for repetitiveness by constructing a 15-mer histogram from the full genome sequence and then calculating the average 15-mer count across each probe, sliding a window size of 15 bases across the length of each probe. Any probe with an average 15-mer count greater than 100 was considered to be repetitive and excluded from further characterization. Non-repetitive probes were then scored for uniqueness by aligning each capture probe to the full cat genome using SSAHA^32^. A close match to the genome was defined as a match length of 30 bases, allowing up to five insertions/deletions/substitutions. Capture probes were selected for each coding sequence feature by scoring one to four probes in a 20-base window, based on repetitiveness, uniqueness, melting temperature and sequence composition, and then choosing the best capture probe in that window. The start of the 20 base windows was then moved 40 bases downstream and the process repeated. Selected probes were allowed to start up to 30 bases before the 5’ start of each feature and overhang the 3’ end by 30 bp. A maximum of five close matches in the genome was allowed when selecting the capture probes.

### Samples and DNA Isolation

Cat DNA samples for WES were donated by owners and archived in accordance with the University of Missouri Institutional Animal Care and Use Committee protocol study protocols 9056, 9178, and 9642. DNA was isolated from 41 whole blood or tissue cat samples using standard organic methods ^33^ and verified for quantity and quality by DNA fluorescence assay (Qubit, Thermo Fisher) and ethidium bromide staining after 0.7% agarose gel electrophoresis. Ten cats with existing whole genome sequence (WGS) data were initially tested followed by 31 novel cats.

### Sequencing

Genomic DNA (250 ng) was fragmented on the Covaris LE220 instrument targeting 250 bp inserts. Automated dual indexed libraries were constructed with the KAPA HTP library prep kit (Roche) on the SciClone NGS platform (Perkin Elmer). The libraries were PCR amplified with KAPA HiFi for 8 cycles. The final libraries were purified with a 1.0x AMPureXP bead cleanup and quantitated on the Caliper GX instrument (Perkin Elmer) and were pooled pre-capture generating a total 5μg library pool. Each library pool was hybridized with a custom Nimblegen probe set (Roche), targeting 35.9 Mb. The libraries were hybridized for 16 - 18 hours at 65°C followed by washing to remove spuriously hybridized library fragments. Enriched library fragments were eluted following isolation with streptavidin-coated magnetic beads and amplified with KAPA HiFi Polymerase prior to sequencing. PCR cycle optimization is performed to prevent over amplification of the libraries. The concentration of each captured library pool was accurately determined through qPCR utilizing the KAPA library Quantification Kit according to the manufacturer’s protocol (Roche) to produce appropriate cluster counts prior to sequencing. The lllumina NovaSeq6000 instrument was used to generate 150 bp length sequences to yield an average of 14 Gb of data per 35.9 Mb target exome, producing ~60x genome coverage. Exome sequencing data are available at the Sequence Read Archive under accession number PRJNA627536.

### Variant Discovery

The following tools/packages were applied to WGS and WES samples in accordance with variant processing as previously described^34^, BWA-MEM version 0.7.17^35^, Picard tools version 2.1.1 (http://broadinstitute.github.io/picard/), Samtools version 1.9,^36^ and Genome Analysis toolkit version 3.8^37,38,39^. Code used for the variant calling workflow can be found at https://github.com/mu-feline-genome/batch_GATK_workflow. For WES processing, GATK tools were restricted to exons annotated in Ensembl release 97 with an additional 100 bp of flanking sequence^40^. Following processing, samples were genotyped in three separate cohorts. The first cohort consisted of all 41 WES samples. The second and third cohorts were ten matched WES and WGS samples. Variants in all three cohorts were tagged using the same variant filtering criteria. For SNVs the filtering criteria was, QD < 2.0, FS > 60.0, SOR > 3.0, ReadPosRankSum < −8.0, MQ < 40.0, and MQRankSum < −12.5. For indels the filtering criteria was, QD < 2.0, FS > 200.0, SOR > 10.0, and ReadPosRankSum < −20.0. Although five Y chromosome genes were included in the exome probe set, these genes had not been added to the aligning reference. For WGS/WES comparison, matched WES/WGS samples were annotated using variant effect predictor (VEP)^41^. Variants from both cohorts were independently tagged as whether they were biallelic, SNPs, or passed filtering criteria. Prior to analysis, variants flanking the exome primary target regions +/− 2bp were removed (**Supplementary Data S1**). Variant processing and comparisons were performed in the R statistical environment using the vcfR package^42^. Common variants between both platforms were determined as those at the same position with the same reference and alternate alleles. Exclusive variants were determined as those where the position and/or the alleles were specific to a particular platform. The initial ten WES cats also had WGS data and are in the SRA under BioProject PRJNA308208 as part of the 99 Lives Cat Genome Sequencing Consortium^29^. Each cat had approximately 30x WGS coverage using an Illumina HiSeq 2500 PE 125 bp using both 350 bp and 550 bp insert libraries.

### Disease and Trait Variant Detection

Variants for all 41 cats were evaluated using VarSeq software (GoldenHelix, Inc.). SNVs were annotated as having high, moderate or low impacts on gene function. High impact variations were those that were a protein truncating variant caused by stop gain or loss and splice-site acceptor or donor mutations^43^. Moderate impacts include missense mutations or frame insertions, and lastly low impact variants are characterized by synonymous base changes, splice region variants or stop retained variance. Known variants for diseases and traits were evaluated in each cat.

### Polycystic Kidney Disease

A pointed cat of the Siberian breed (a.k.a. Neva Masquerade) was diagnosed with polycystic kidney disease based on signs of renal disease (polydipsia, polyuria) and ultrasonography. DNA was submitted using buccal swabs and a whole blood sample to two different commercial testing laboratories in which both confirmed the absence of the currently known autosomal dominant polycystic kidney disease in *polycystin-1 (PKD1)^44,45^*. The dam and a sibling were also confirmed as having PKD by ultrasonography but we not available for genetic analyses.

### Cystinuria

A three-month-old European shorthair kitten from the isle of Korfu, Greece, was presented to the AniCura Small Animal Hospital, Bielefeld, FRG, for heavy straining during urination and the owner report the kitten would fall over from time to time. The kitten had been pretreated with two injections of cephalexine and dexamethasone for suspected cystitis, however, difficulty in urination worsened. Upon hospital admission, the kitten was in good general condition. Abdominal palpation revealed an enlarged urinary bladder. Abdominal X-ray showed over 30 radiolucent urinary stones up to a diameter of half of the width of the last rib. Urinary bladder stones and some urethral stones were removed via cystolithotomy and retrograde flushing of the urethra. Urinary stones were submitted for infraspectroscopic stone analysis. Stone analysis revealed pure cystine stones and a diagnosis of cystinuria was made. Urinary stones reoccurred at six months of age, but they kitten was otherwise healthy.

## Results

### Phenotype cohort

The 41 cats in exome study represent different diseases and traits, some with known disease alleles others unknown (**Table 1**). The initial ten cats had nine known disease variants and various known mutations for coat colors and fur types. In the group of 31 novel exomes, the cats represented 11 different breeds and 14 random bred cats. Seven pairs of cats were sequenced to evaluate causes for mediastinal lymphoma, a seizure disorder, eyelid colobomas, hypothyroidism, hypovitaminosis D, blue eyes of Ojos Azules breed, and curly hair coat of the Tennessee Rex. Five cats were reported with cardiac diseases, including hypertrophic cardiomyopathy (HCM). At least seven neurological disorders are represented in the study population, generally representing novel presentations in random bred cats. Overall, the 41 cats had approximately 31 different unknown disease presentations.

**Table 1.** Signalment and diseases of 41 cats for WES evaluation.

\*Mutations as tentative causal variants for diseases presented.
| No. | Id. | Breed | Sex | Disease / Trait | Gene(s) |
| --- | --- | --- | --- | --- | --- |
| 1 | 19725 | Lykoi | F | Lykoi | *HR* |
| 2 | 13230 | Mixed Breed | F | Bengal PRA / Bobbed tail | *KIF3B / HES7* |
| 3 | 14056 | Mixed Breed | M | Persian PRA / *Long* | *AIPL1 / FGF5* |
| 4 | 17994 | Mixed Breed | F | Hydrocephalus | *GDF7* |
| 5 | 19067 | Munchkin | F | Dwarfism / Dominant White | *UGDH / KIT* |
| 6 | 5012 | Oriental | M | Lymphoma | *Unknown* |
| 7 | 20382 | Peterbald | M | *Hairless* | *LPAR6\** |
| 8 | 11615 | Random Bred | M | *Dominant White* | *KIT* |
| 9 | 18528 | Random Bred | M | *Spotting* | *KIT* |
| 10 | 20424 | Siberian | F | *Long* / Cardiac disease | *FGF5 / Candidate* |
| 11 | 22550 | Bengal | F | Polyneuropathy | *Unknown* |
| 12 | 20957 | Devon Rex | U | Papilloma virus | *Unknown* |
| 13 | 22752 | Devon Rex | M | Neurological disorder | *Unknown* |
|  | 21464 |  |  |  |  |
| 14 -15 | 21983 | Ojos Azules | 1F:1M | Ojos Azules | *Unknown* |
| 16 | 20964 | Oriental | F | Cardiac disease | *Unknown* |
| 17 | 22728 | Random bred | F | Cystinuria | *SLC3A1\** |
| 18 | 20617 | Random Bred | M | Neuronal ceroid lipofuscinosis | *Candidate* |
| 19 | 20948 | Random Bred | M | Cinnamic acid urea | *Unknown* |
| 20 | 21153 | Random Bred | M | Ambulatory paraparesis | *Unknown* |
| 21 | 22287 | Random Bred | F | Myotonia congenita | *Unknown* |
| 22 | 22397 | Random Bred | M | Neurological disorder | *Unknown* |
| 23 | 22505 | Random Bred | M | Cardiac disease | *Unknown* |
| 24 | 22623 | Random Bred | U | Pycnodysostosis | *Candidate* |
| 25 | 22740 | Random Bred | F | Epidemolysis bullosa | *Unknown* |
| 26 – 27 | 22741 | Random Bred | F:M | Eyelid coloboma | *Unknown* |
| 28 | 22751 | Random Bred | M | Ehlers-Danlos | *Unknown* |
| 29 – 30 | 22763 | Random Bred | 2F | Hypothyroidism | *Candidate* |
| 31 – 32 | 22761 | Savannah | 2M | Hypovitaminosis D | *Unknown* |
| 33 | 21984 | Scottish Fold | F | Cardiac disease | *Candidate* |
| 34 – 35 | 20384 | Selkirk Rex | 1F:1U | Seizures | *Unknown* |
| 36 | 20953 | Siamese | F | Cardiac disease | *Candidate* |
| 37 | 22622 | Siberian | U | PKD | *PKD2\** |
| 38 | 22711 | Singapura | F | Hypovitaminosis D | *Candidate* |
| 39 – 40 | 8641 | Tennessee Rex | 1F:1M | Rexoid hair coat | *Unknown* |
| 41 | 6623 | Oriental | M | Lymphoma | *Unknown* |
| **41** |  | **14 breeds** | **19F:18M:4U** | **~31 diseases & traits** |  |

### Sequence coverage and specificity

To assess the performance of the feline WES resource, WES data was produced on ten cats that had WGS data for comparison. Approximately 55 – 259 million raw reads generated per sample (**Supplementary Table 1, Supplementary File 2**). After mapping to Felis_catus_9.0, base quality trimming, and duplicates removal, the percentage of unique reads that mapped to the cat genome assembly was ~82%. The average sequencing depth was 267x with a range of 76x to 458x (**Supplementary Table 2, Supplementary File 2**). Of the 201,683 exonic targets, 98.1% of the exonic sequences had aligned coverage >20x with an average of 6.47% off-target sequence (**Supplementary Table 3, Supplementary File 2).** For the novel 31 cat exomes, the average coverage was 80x ranging from 60 – 108x. The percent of on target coverage up to 10x was generally 98 – 99%. The percent of on target based covered >20x was 96.41%, ranging from 91 – 98% with an average of 10.41% off-target sequence. Across all cats, an average of 82% of reads aligned with a range of 75% to 85% with an average of >99% of the bases aligned. When looking at targeted bases, an average of 99% of bases aligned with at least 2x coverage. There was a reduction at deeper coverage, for example at 40 and 100x, 93.5 and 58% of targeted bases were covered, respectively (**Figure 1**). Approximately 70M reads produced approximately 80x coverage, which generally ensured >98% of bases had 20x coverage.

**Figure 1.**
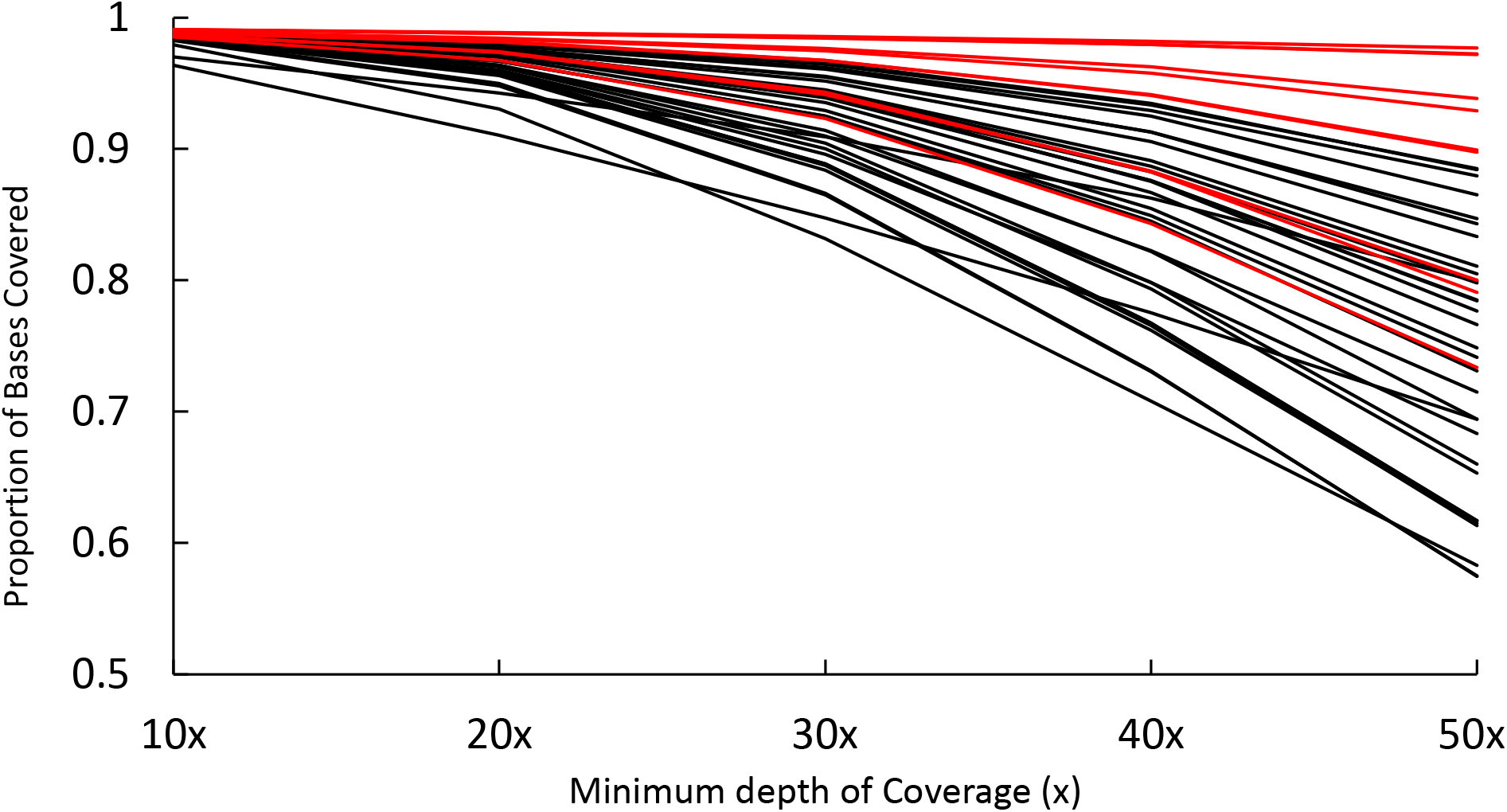
The proportion of bases covered with the exome capture probes. The initial 10 ples are colored in red, with the X axis showing the depth of coverage, which is how many es a nucleotide base is covered starting at a depth of 10x and increasing to 50x.

### WGS versus WES-specific variant discovery

A set of common variants and platform (WGS versus WES) exclusive variants were defined and then filtered for quality, variant type, and biallelic status. For high impact variants, WES and WGS identified 582 and 617 SNPs, respectively, with 97.8% of the WES SNPs also identified by WGS and 92.1% of the SNPs also identified by WES (**Table 2**). The most exclusive percentage of identified variants were for splice donor / acceptor sites and stop gains, however, the overall count of these variants was low, ranging from 3 to 19 total variants. Moderate (missense) and low impact variants had very high concordance between the WES and WGS datasets, ranging from 94.7% for 3’ UTR SNPs in WGS to ~100% for most SNPs identified by WES. Altogether only a small fraction of SNPs (WES = 834 and WGS = 2,195) were exclusive to a particular platform (**Figure 2a**). Considering small indels, the WES and WGS data had lower concordance than SNPs (**Table 3**). Although WES detected 1,738 high impact indels and WGS detected 1,931, the percentage of commonly identified and exclusive indels showed more variation between consequence categories than SNPs. For both SNPs and indels, high impact mutations represented a disproportionate fraction of the platform exclusive variants.

**Figure 2:**
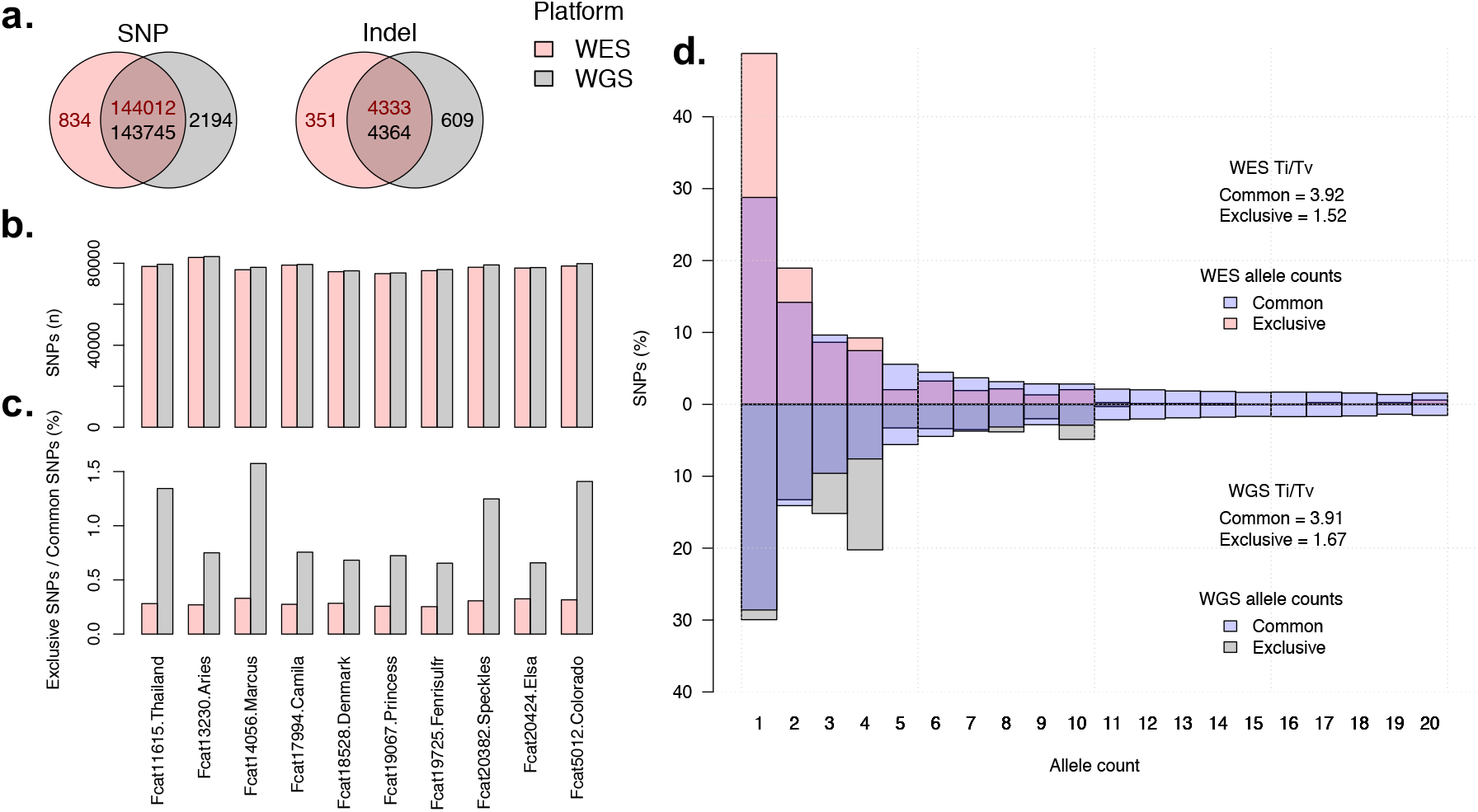
Variant calling statistics for 10 cats sequenced on both platforms. **a**) Venn diagrams showing the number of exclusive and common variants per platform. Dark red text indicates the number of variants found in WES and black text indicates the number of variants found in WGS. The reason the number of common variants differ between platforms is because common variants were identified prior to filtering. **b**) The number of SNPs found in each sample in both platforms. **c**) The percentage of SNPs found as exclusive to each sample for each platform. The first, third, eighth, and tenth samples are males. All other samples are female. **d**) Allele count distribution for common and exclusive SNPs in both platforms. WES SNPs are shown on top and WGS SNPs are shown upside down on the bottom. In addition, the Ti/Tv ratio for sets of SNPs is also shown

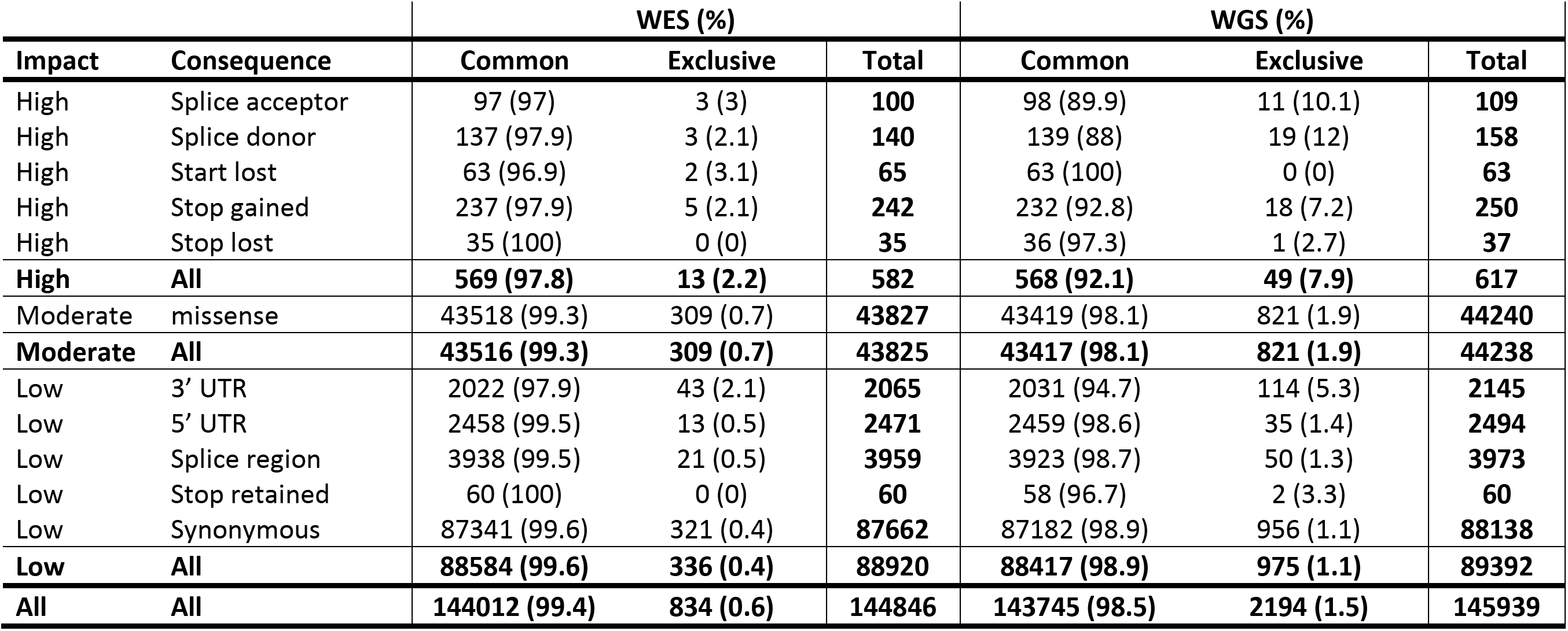

|  |  | WES (%) |  |  | WGS (%) |  |  |
| --- | --- | --- | --- | --- | --- | --- | --- |
| Impact | Consequence | Common | Exclusive | Total | Common | Exclusive | Total |
| High | Splice acceptor | 97 (97) | 3 (3) | **100** | 98 (89.9) | 11 (10.1) | **109** |
| High | Splice donor | 137 (97.9) | 3 (2.1) | **140** | 139 (88) | 19 (12) | **158** |
| High | Start lost | 63 (96.9) | 2 (3.1) | **65** | 63 (100) | 0 (0) | **63** |
| High | Stop gained | 237 (97.9) | 5 (2.1) | **242** | 232 (92.8) | 18 (7.2) | **250** |
| High | Stop lost | 35 (100) | 0 (0) | **35** | 36 (97.3) | 1 (2.7) | **37** |
| **High** | **All** | **569 (97.8)** | **13 (2.2)** | **582** | **568 (92.1)** | **49 (7.9)** | **617** |
| Moderate | missense | 43518 (99.3) | 309 (0.7) | **43827** | 43419 (98.1) | 821 (1.9) | **44240** |
| **Moderate** | **All** | **43516 (99.3)** | **309 (0.7)** | **43825** | **43417 (98.1)** | **821 (1.9)** | **44238** |
| Low | 3' UTR | 2022 (97.9) | 43 (2.1) | **2065** | 2031 (94.7) | 114 (5.3) | **2145** |
| Low | 5' UTR | 2458 (99.5) | 13 (0.5) | **2471** | 2459 (98.6) | 35 (1.4) | **2494** |
| Low | Splice region | 3938 (99.5) | 21 (0.5) | **3959** | 3923 (98.7) | 50 (1.3) | **3973** |
| Low | Stop retained | 60 (100) | 0 (0) | **60** | 58 (96.7) | 2 (3.3) | **60** |
| Low | Synonymous | 87341 (99.6) | 321 (0.4) | **87662** | 87182 (98.9) | 956 (1.1) | **88138** |
| **Low** | **All** | **88584 (99.6)** | **336 (0.4)** | **88920** | **88417 (98.9)** | **975 (1.1)** | **89392** |
| **All** | **All** | **144012 (99.4)** | **834 (0.6)** | **144846** | **143745 (98.5)** | **2194 (1.5)** | **145939** |

**Table 3:**
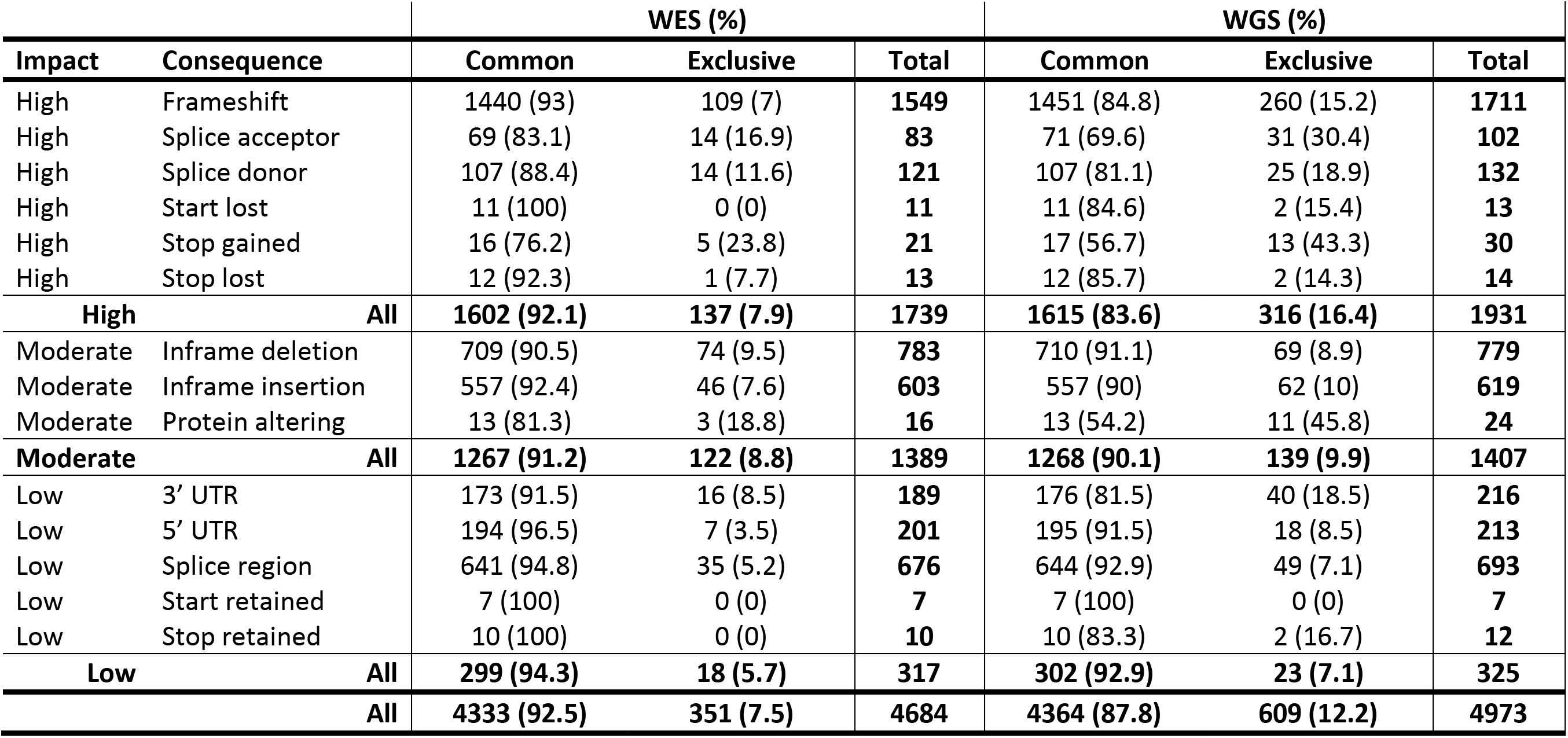
Indel consequence counts of WES versus WGS as determined by variant effect predictor.

|  |  | WES (%) |  |  | WGS (%) |  |  |
| --- | --- | --- | --- | --- | --- | --- | --- |
| **Impact** | **Consequence** | **Common** | **Exclusive** | **Total** | **Common** | **Exclusive** | **Total** |
| High | Frameshift | 1440 (93) | 109 (7) | **1549** | 1451 (84.8) | 260 (15.2) | **1711** |
| High | Splice acceptor | 69 (83.1) | 14 (16.9) | **83** | 71 (69.6) | 31 (30.4) | **102** |
| High | Splice donor | 107 (88.4) | 14 (11.6) | **121** | 107 (81.1) | 25 (18.9) | **132** |
| High | Start lost | 11 (100) | 0 (0) | **11** | 11 (84.6) | 2 (15.4) | **13** |
| High | Stop gained | 16 (76.2) | 5 (23.8) | **21** | 17 (56.7) | 13 (43.3) | **30** |
| High | Stop lost | 12 (92.3) | 1 (7.7) | **13** | 12 (85.7) | 2 (14.3) | **14** |
| **High** | **All** | **1602 (92.1)** | **137 (7.9)** | **1739** | **1615 (83.6)** | **316 (16.4)** | **1931** |
| Moderate | Inframe deletion | 709 (90.5) | 74 (9.5) | **783** | 710 (91.1) | 69 (8.9) | **779** |
| Moderate | Inframe insertion | 557 (92.4) | 46 (7.6) | **603** | 557 (90) | 62 (10) | **619** |
| Moderate | Protein altering | 13 (81.3) | 3 (18.8) | **16** | 13 (54.2) | 11 (45.8) | **24** |
| **Moderate** | **All** | **1267 (91.2)** | **122 (8.8)** | **1389** | **1268 (90.1)** | **139 (9.9)** | **1407** |
| Low | 3' UTR | 173 (91.5) | 16 (8.5) | **189** | 176 (81.5) | 40 (18.5) | **216** |
| Low | 5' UTR | 194 (96.5) | 7 (3.5) | **201** | 195 (91.5) | 18 (8.5) | **213** |
| Low | Splice region | 641 (94.8) | 35 (5.2) | **676** | 644 (92.9) | 49 (7.1) | **693** |
| Low | Start retained | 7 (100) | 0 (0) | **7** | 7 (100) | 0 (0) | **7** |
| Low | Stop retained | 10 (100) | 0 (0) | **10** | 10 (83.3) | 2 (16.7) | **12** |
| **Low** | **All** | **299 (94.3)** | **18 (5.7)** | **317** | **302 (92.9)** | **23 (7.1)** | **325** |
|  | **All** | **4333 (92.5)** | **351 (7.5)** | **4684** | **4364 (87.8)** | **609 (12.2)** | **4973** |

Across samples, each individual cat carried approximately 80,000 SNPs total, with only marginal differences between platforms and individuals (**Figure 2b**). Alternatively, platform exclusive SNPs, particularly for WGS, did not exhibit these same patterns. The four male cats, each carried approximately twice as many WGS exclusive variants as female cats (**Figure 2c**).

Another method for characterizing platform exclusive SNPs, is to measure their allele count distributions. Compared to common SNPs identified from WES, exclusive SNPs were heavily skewed toward allele counts of one (**Figure 2d**). Using common SNPs as a truth set for comparison, the WES exclusive allele count distribution is consistent with SNPs identified by random error, as most of these SNPs only appear once in the dataset. Moreover, this result is reflected by the Ti/Tv ratios of each dataset. WES common SNPs are at 3.92, indicating a low concentration of false positive variant sites, while WES exclusive SNPs are at 1.52, indicating a high concentration of false positive variant sites. Alternatively, allele counts for WGS exclusive SNPs have two peaks. The first is at an allele count of one, which is similar to WES exclusive SNPs, and the second is at an allele count of four, which is suggestive of more systematic error in variant detection. This second peak for WGS exclusive SNPs is likely consistent with the increased WGS exclusive variant detection observed in male cats. For WGS SNPs, the Ti/Tv ratios for both common and exclusive SNPs is similar to WES SNPs, where exclusive SNPs are enriched for false positive variant sites.

### WGS versus WES bias in variant discovery

To detect bias toward specific genes using the WGS and WES platforms, the number of variants per gene was compared between WGS and WES results (**Supplementary Data S2**). A large number of genes had greater than 20 more WGS variants than WES variants (**Figure 3**). To investigate the cause for these outliers, the top 50 of these outlier genes were selected for further analysis (**Supplementary Data S3**). Of these, 14 genes were found on the X chromosome, suggesting these differences in variant detection may correspond to the increased number of WGS exclusive SNPs in males observed in (**Figure 2c) (Supplementary data S3).** Apart from enrichment on chromosome X, another cluster of 13 genes with WGS biased variant detection were located on chromosome D1. These genes were mostly olfactory receptors, which are usually repetitive and therefore difficult to design unique bait probes. Another gene of note, LOC101099449, contained 713,328 bp of target sequence. When analyzed more closely, LOC101099449’s target sequence overlapped an entire Immunoglobulin lambda locus at chromosome D3:20097014 - 20810341, a region that is usually highly variable between individuals. All other genes with WGS-biased variant detection were distributed randomly.

**Figure. 3.**
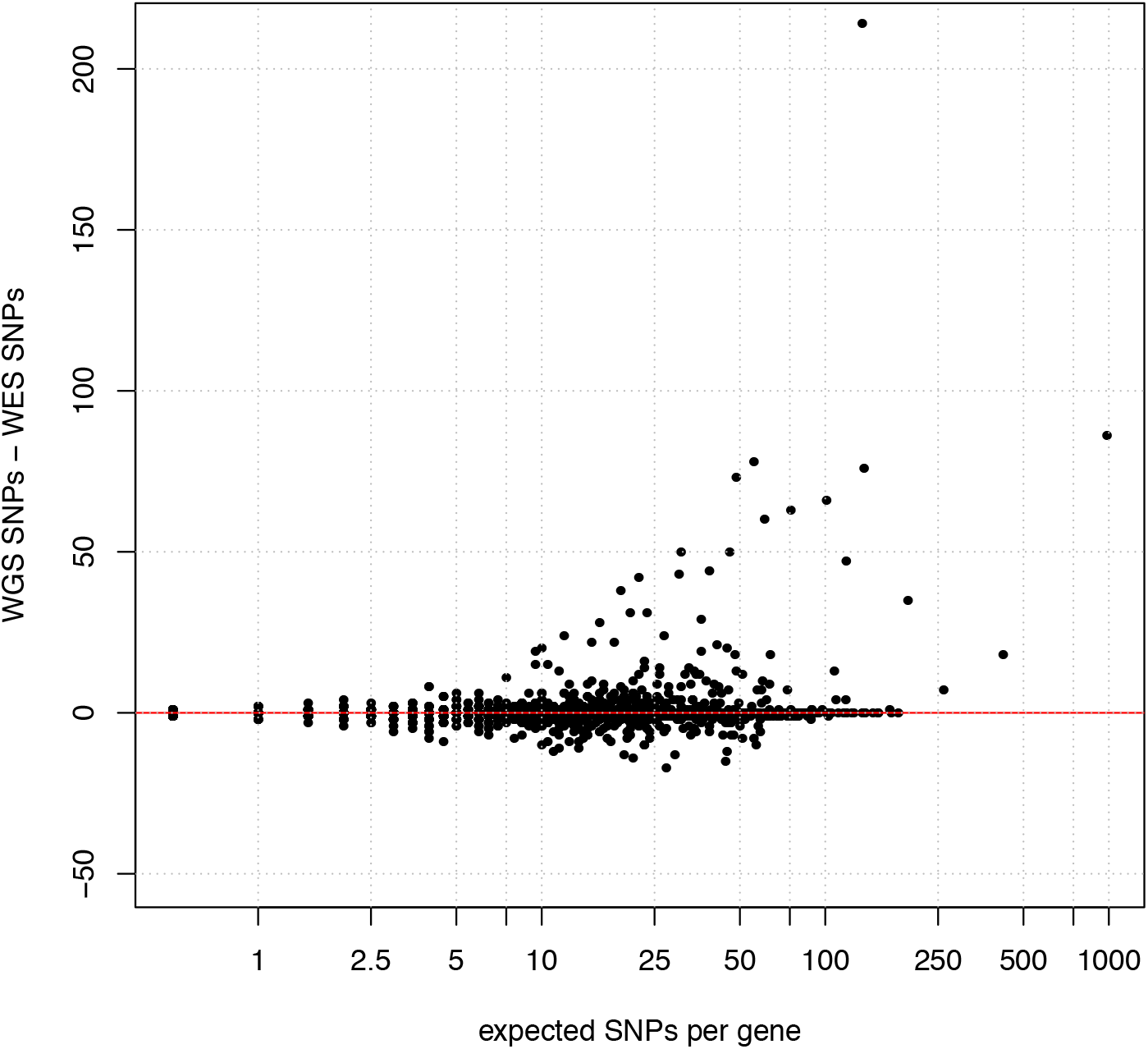
Gene-wise platform bias. Each individual point on the scatterplot is a gene with the y axis displaying differences in SNP counts per gene. Genes with more WGS SNPs than WES SNPs have positive values, where genes have negative values when there is more WES SNPs instead. Expected SNP number is calculated as the mean number of SNPs per gene across both platforms and is plotted on a log scale.

To further investigate increased WGS-biased variant detection on chromosome X, the mean number of variants per individual was compared between males and females (**Table 4**). Across autosomes and sequencing platforms, sex-based percentage differences were relatively low, ranging between 7% and 10%. Alternatively, across both gene groupings, the percentage difference between the sexes on the X chromosome were much higher. For the top 50 WGS outlier genes, both platforms showed an approximate 98% sex difference, whereas all genes showed a 61.09% sex difference for WGS and a 45.02% sex difference for WES. Since the percentage sex difference in outlier genes is similar across both platforms, results suggest that platform bias on chromosome X is more likely due to platform specific increased variant detection in these regions, rather than differential abilities of platforms to detect variants in either sex. Importantly, the actual number of chromosome X sex differences in both platforms is similar across gene groupings. In the top 50 WGS outliers, the difference between the chromosome X mean male and female SNP counts is 1340.92, while across all X chromosome genes this same difference is equal to 1202.5 (**Table 4**).

**Table 4:** Mean SNPs per individual for ten WES and WGS cats.

| **Genes** | **Top 50 WGS outliers** |  |  |  |  |  |
| --- | --- | --- | --- | --- | --- | --- |
| **Platform** | **WGS** |  |  | **WES** |  |  |
| **Sex** | Male | Female | Difference (%)<sup>1</sup> | Male | Female | Difference (%)<sup>1</sup> |
| Autosome | 1595.00 | 1445.67 | **149.33 (9.36)** | 946.25 | 872.83 | **73.42 (7.76)** |
| X chromosome | 1363.75 | 22.83 | **1340.92 (98.33)** | 829.75 | 23.00 | **806.75 (97.73)** |
| **Genes** | **All** |  |  |  |  |  |
| **Platform** | **WGS** |  |  | **WES** |  |  |
| **Sex** | Male | Female | Difference (%)<sup>1</sup> | Male | Female | Difference (%)<sup>1</sup> |
| Autosome | 53724.75 | 57605.50 | **3880.75 (7.22)** | 53189.50 | 57217.67 | **4028.17 (7.57)** |
| X chromosome | 1968.50 | 766.00 | **1202.50 (61.09)** | 1412.00 | 776.33 | **635.67 (45.02)** |

<sup>1</sup>Percentage differences in parentheses were calculated as a fraction of mean SNPs per male individual.

To examine the potential overlap between platform and sex bias, the distribution of SNPs per gene along chromosome X were analyzed. Platform biased genes are clustered between positions 15 to 70 Mb (**Figure 4a**). Across both platforms, these genes also have the highest SNP concentration, with > 20 SNPs per kb of coding sequence (**Figure 4a**).

**Figure 4:**
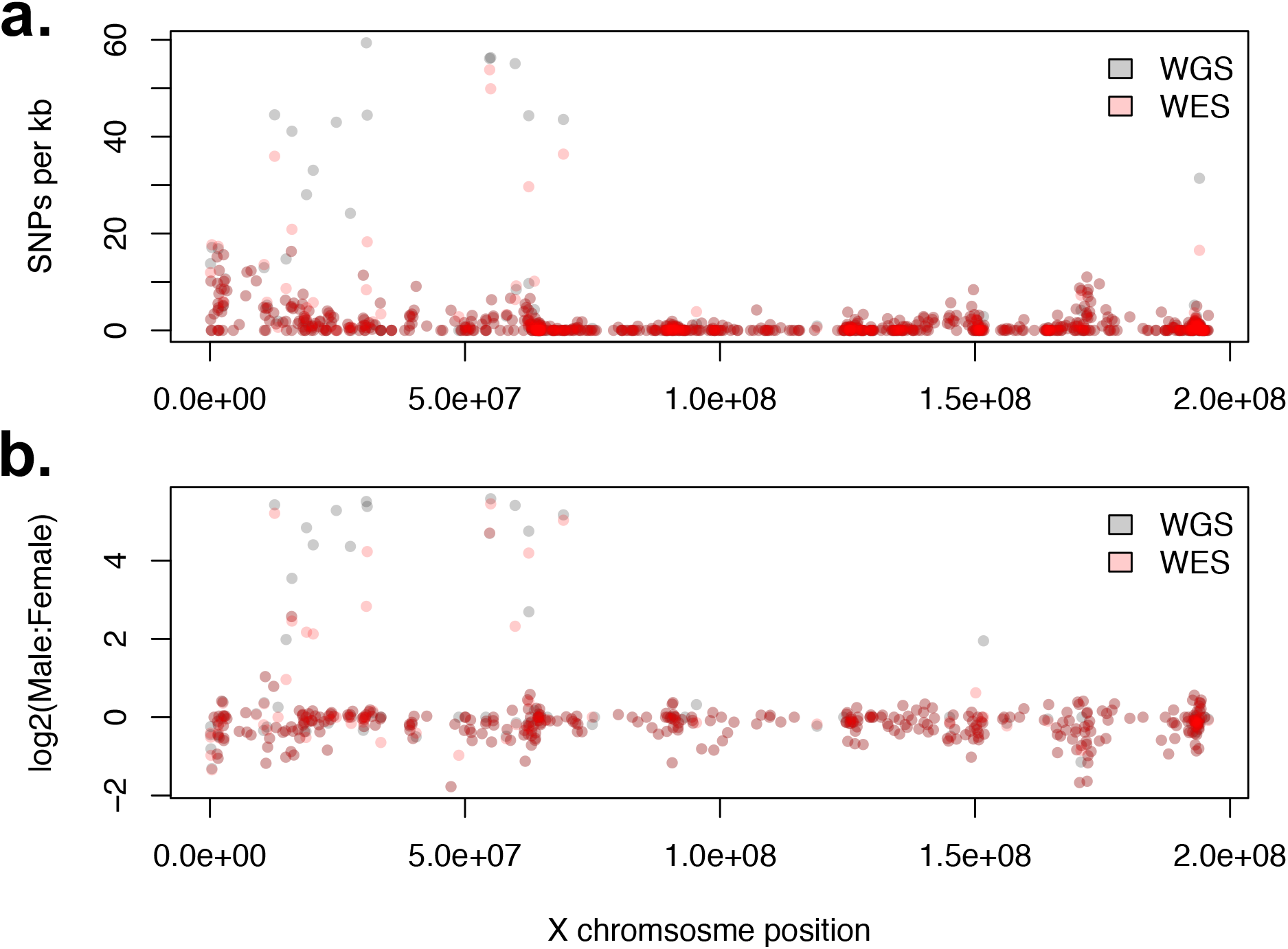
Distribution of SNPs per gene along chromosome X. **a**) Total SNPs per kb of coding sequence per gene. **b**) Sex biased variant detection along chromosome X. Bias is calculated as fold change ratio between the mean number of SNPs per individual per gene for males and females. Specifically, this was calculated for each gene as log2((mean male SNPs + 1) / (mean female SNPs + 1)). The ones were added to remove undefined results caused by dividing by the number 0.

Alternatively, the majority of genes outside this region have SNP concentrations of < 5 SNPs per kb of coding sequence. Regarding sex bias, while the overall percentage difference across platforms is similar (**Table 4**), individual genes show platform specific variability in effect size. A larger number of WGS genes than WES genes show a 4-fold bias toward variant detection in males (**Figure 4b**). However, despite this variation across platforms, the genes with increased sex bias are indeed the same genes with increased platform bias (**Supplementary Data S4**). Therefore, on chromosome X, platform biases and sex biases in SNP discovery appear confounded, as numerous factors within the same genes are relatively consistent across both platforms. This suggests both biases have a similar underlying root cause differently expressed in each platform.

A potential cause of sex bias in variant discovery is that the biased genes have degraded copies on the Y chromosome. For the ten known feline X chromosome genes with degraded Y copies,^46^ the total number of SNPs per platform and the mean number of SNPs per individual were calculated. Of these ten genes, nine have platform specific differences in SNP discovery greater than 11 and are therefore among the top 50 outlier genes for platform specific bias (**Supplementary Table 4**). Moreover, almost all SNPs found in these genes were found only in males, regardless of platform. For WGS there was an average total of 1169.25 SNVs found in males with only an average total of 7.83 found in females. For WES the numbers were similar with an average total of 774.5 SNPs found in males and an average total of 7.83 SNPs found in females (**Supplementary Table 4**). Together these results indicate a major portion of sex bias in variant discovery is due to the absence of a Y chromosome in the Felis_catus_9.0 assembly.

### Known variant validation

Using Ensembl 99 for annotation with selection of exons with +/− 30 bp to match exome capture design and visualized using VarSeq (GoldenHelix, Inc). A majority of the 115 variants in the domestic cat documented as causal for diseases and traits affect the coding regions or a splice donor/acceptor site^47^. Forty-four known variants were identified in the WES cohort. All variants for coat colors and diseases known to be present in the ten cats were identified, including the alleles in the loci for *Agouti* (*ASIP - a ^48^), Brown (TYRP1 – b)^49^, Color (TYR – c^s^)^50^, Dense (MLPH)^51^*, *Longhair* (*FGF5*)^52^, Lykoi (*HR*)^53^, Bengal (*KIF3B*)^54^ and Persian progressive retinal degeneration (*AIPL1*)^27^, hydrocephalus (*GDF7*)^55^, and others (**Supplementary Data 5)**. The cats had various known mutations affecting cat blood type. As anticipated, the *KIT* intron 1 structural variants for *White* and *Spotting* were not identified, as well as the structural variant in *UGDH* causing dwarfism^27^.

### Novel candidate variant discovery

Novel DNA variants were explored as causal for diseases and traits in 33 cats. A novel frameshift mutation in *polycystin 2* (*PKD2)^56^*, a gene associated with PKD, was predicted to disrupt protein function in a Siberian cat shown by ultrasound to have PKD. The c.2211delG causes a p.Lys737Asnfs*2 at position B1:134992553. This variant was heterozygous in the affected cat and unique to the exome data and not identified in the 195 cat 99 Lives variant dataset^29^.

The *lysophosphatidic acid receptor 6 (LPAR6)* c.250_253delTTTG variant that causes a p.Phe84Glufs*9 and is associated with the autosomal recessive rexoid (marsella wave) coat of the Cornish rex breed was detected in a Peterbald cat, which is a hairless breed ^57,53^. However, the hairless trait is considered autosomal dominant by cat breeders. The annotation also suggested a c.249delG causing a p.Phe84Leufs*10, therefore, this Peterbald cat is suggested as a compound heterozygous for two mutations juxtaposed in *LPAR6*. This variant was heterozygous in the affected cat and unique to the exome data and not identified in the 195 cat 99 Lives variant dataset. Known feline disease variants were also re-identified (**Supplementary Data 5)**^29^. A *solute carrier family 3 member 1* (*SLC3A1)* variant was homozygous in a Greek cat presenting with cystinuria^58^. The c.1342C>T causing a p.Arg448Trp at position A3:66539609 has been previously documented to be associated with this condition. No other cat in the exome dataset had this variant. Many of the variants associated with cat blood group B and its extended haplotype were detected in one to 11 cats, suggesting five cats as Type B, one was confirmed^59^. Variants were detected in *APOBEC3*, which is associated with FIV infection in cats, and three cats had the allelic combination that produces the IRAVP amino acid haplotype that is associated with FIV resistance^60^. Unexpectedly, two cats were heterozygous for a porphyria variant in *UROS* (c.140C>T, c.331G>A)^61,62^, one cat was homozygous for *FXII* deficiency variant (FXII_1631G>C)^61^, which had died as a kitten, and one cat was heterozygous for a copper metabolism deficiency in *ATP7B^63^*. Additional variants for neuronal ceroid lipofuscinosis, pycnodysostosis, Ehlers-Danlos syndrome, hypothyroidism, and hypovitaminosis D, and several individual specific variants for hypertrophic cardiomyopathy are under further investigation (**Table 1**).

## Discussion

Whole exome sequencing has flourished over the past decade and is becoming the state-of-the-art technology for Precision / Genomic Medicine. Well recognized for clinical applications in human medicine, the success of WES is highly dependent on the accuracy of the genome assembly and annotation^10^. Well annotated genomes, such as human and mice, has allowed the development of various exome capture products that range from particular genes of focus for clinical applications to more extensive designs that include 5’ and 3’ untranslated regions, miRNA, lncRNA, suspected regulatory elements and variation is the exon flanking sequence length. For mammals with ~2.4 – 3.0 Gb genomes, exome designs have included 49.6 Mb for mouse, 54 Mb for cow, 71 Mb and 146.8 Mb in rats^64,65^ and ~48.2 Mb for humans. In domestic dog, products have ranged from ~53 Mb – 152 Mb, capturing up to 6% of the genome, with an overlap of ~34.5 Mb of the genome between the capture designs^15-18^. Overall, the size of the capture design is a balance between sequencing costs, larger designs imply higher costs, and the intended applications of the product. Presented is a WES capture platform designed specifically for *Felis Catus*.

The sequence capture probes for the cat WES were designed from the annotated Felis_catus_9.0 genome assembly, which is one of the more robust long-read based assemblies for mammals, strongly supporting efficient design^29^. The design included 35 Mb, primarily focusing on exomes and the flanking regions to detect splice door – acceptor variants as little annotation for miRNA is available for cats. The total gene count in cats is slightly smaller as compared to dogs, with dogs having 291 more protein coding sequences, contributing to a smaller target size in cat. The success of disease variant identification is dependent on several factors, including sequencing depth and efficient design of the probes that allow adequate read coverage for variant detection. The success of disease variant identification is dependent on several factors, including sequencing depth, and efficient design of the probes that are on target, thereby reducing waste in sequencing costs, and allow adequate read coverage for variant detection. The percent of unique reads was consistent for all cats, averaging 81 – 82%, and nearly 100% aligned to the cat genome as intended. A read coverage of ~20x is regarded as the standard to efficiently detect heterozygous variants ^66^. In the first 10 cats sequenced, the mean 267x coverage indicates a maximum coverage of 99% of the exonic sequences had aligned coverage >20x coverage. For the 31 cats with an average coverage of 80x, 96.41% of the bases were on target with greater than 20x coverage. In comparison to the first domestic dog exome design, which covered 52.8 Mb (<2% of the genome) divided over 203,059 regions, at a lower mean sequencing depth over 8 samples (102x), the dog design had a higher percentage of mapped reads at ~87 – 90%. However, when comparing base coverages, 93 to 94% (<9 Mb) of the targeted bases (<53 Mb) were covered at least once and 89 to 91% were covered at least five times in the canine design^15^, while the cat coverages were higher at nearly 100%. The pig exome capture probes demonstrate 90% of bases covered at 20x coverage, discovering 264,000 SNPs and indels^67^. Overall, direct comparisons are difficult due to the differences in annotation, genome assembly accuracy, and design techniques. For example, the cat design included 30 bp flanking the exon boundaries and a maximum of five close matches in the genome was allowed when selecting the capture probes. Both of these attributes were zero for the canine design.

The intended application of the WES design for the cat is the identification of heritable, Mendelian diseases and phenotypes. To assess the efficiency of the feline exome design, ten matched samples with WGS and WES data were compared. The ten cats had an average of ~30x WGS coverage and ~267x for the WES coverage. The vast majority of SNPs and indels in target regions were detected by both platforms. Altogether, SNP discovery with the feline exome probes was extremely consistent with variant discovery from WGS, 99.4% of WES SNPs were detected in WGS while only 1.5% of WGS SNPs were absent from the WES dataset. Alternatively, indel discovery was less consistent across platforms, where 92.5 % of WES indels were detected in the WGS indel set and 12.2% of WGS indels were absent from the WES indel set. Generally, indel identification is more prone to errors than SNP identification, therefore the reduced indel consistency across platforms may be reflective of their difficulty to correctly identify using either platform. Differences in the number of common variants between platforms is due to differential filtering, as common variants were identified prior to when filtering was performed. However, since high impact mutations are generally rare due to their impact on disease processes, their enrichment within platform exclusive variant sets could be indicative of random errors. In the same manner, low impact variants represent a lower than expected fraction of platform exclusive variants.

For a small number of genes, a larger number of SNPs were detected using WGS. These genes were mostly restricted to olfactory receptors on chromosome D1 and genes on the X chromosome that have degraded copies on the Y. The repetitive nature of olfactory receptors means they are likely to cause complications in hybridization and mapping. Since olfactory receptors are rarely involved in disease, loss of these genes is barely an impediment for diagnostic purposes. For X chromosome WGS biased genes, there was also bias toward increased WGS variant discovery in males. One potential cause is these genes belong to the degenerate X region of the Y chromosome. A collection of 10 known X chromosome genes with degrading Y chromosome copies all showed high levels of sex bias and platform bias^46^. The reason these genes had more variants in males is because the Y chromosome copies contained a large number of mismatches. Similarly, the increased number of variants may have also affected hybridization of Y chromosome fragments to X chromosome probes, leading to reduced detection of variants in WES. Moreover, the number of variants in females for these genes was largely consistent across platforms, indicating that discrepancies are most likely due to the presence of the Y chromosome. The impact from degraded X genes on the Y chromosomes propagated throughout the analysis. WGS exclusive SNPs were more common in males and the allele count distribution contained a peak at an allele count of four. Even though the effect was found across both platforms, it was especially observable in the WGS exclusive dataset and may have otherwise remained hidden. Importantly, while the feline exome set contained probes for *DDX3Y*, *USP9Y*, *UBE1Y*, and *KDM5D*, which are all Y chromosome degraded X genes, these genes were not included in the reference genome used to align reads. Despite, this absence of the partial Y assembly, many Y chromosome degraded X genes do not have probes designed. Overall, both WGS and WES analysis in the cat will be greatly improved by the assembly of a domestic cat Y chromosome, indicating the importance of developing an improved Y chromosome assembly in the cat.

Variants were investigated to identify novel candidate *de novo* mutations. Various known diseases and phenotypes were first confirmed to test the accuracy of the design. Known causative alleles in *Agouti, Brown, Color, Dense, Gloves, Dilution, Extension*, *Long, Lykoi,* and hairless coat types were confirmed^47^. Disease variants were confirmed for candidate alleles in hydrocephalus, hypertrophic cardiomyopathy, and progressive retinal atrophy. However, known structural variants were not detected nor intronic variants, as expected. When analyzing discordant reads in WGS dwarf sample, a deletion and rearrangement indicating a structural variant is visible in the *UDGH* gene. The discordant reads associated with this variant do not show up in the WES analysis (**Supplementary Figure 1**). Therefore, the WES approaches will likely fail to identify structural variants, an important limitation of the designs as SVs may account for up to 50% or more of disease variants^13^.

Novel causal variants are also suggested from the collection of cats used for the WES design study. Feline PKD is a common inherited autosomal dominate disease affecting about 6% of the world’s cats^44^. PKD is characterized by fluid filled cysts than form in in the bilateral kidneys and may even form in the in the liver and pancreas and may lead to renal failure^68^. Many of the features of PKD are similar to human ADPKD and recent studies have cat have demonstrated the utility of the model^24,69^. The c. 10063C>A mutation in exon 29 of *PKD1* was the only known causative allele for cat ADPKD^44^, however, for human ADPKD, variants are found throughout *PKD1*. The *polycystin 2* (*PKD2)* c.2211delG at position B1:134992553 causes a p.Lys737Asnfs*2 and was identified in a Siberian cat from Europe, indicating additional alleles may be segregating for PKD in cats.

Domestic cats have various forms of atrichia and hypotricha that have led to the development of specific breeds. The two breeds that are recognized as completely hairless are the Sphynx and Donskoy. Donskoy cats are a breed of Russian cats where the loss of hair is determined by semi-dominant gene. Peterbald cats were bred in Russia in 1994 as a product of a Donskoy and an Oriental Shorthair cross, and are often born with no hair, or lose their hair over time. Cornish Rex, a hyprotichia breed, that is characterized by a curly coat, is caused by a homozygous mutation in *LPAR6* ^70^. In this study, the examined Peterbald cat had the 4 base pair deletion *LPAR*6 for the Cornish rex. This cat may be a compound heterozygote for a deletion that is in juxtaposition to the Cornish rex variant. Both variants cause premature stop codons a few amino acids downstream. Several other diseases are under investigation with functional studies to support their causality in diseases.

A variety of additional disease-associated variants were identified in the exome data, including variants for blood types and re-identification of alleles for cystinuria, in which the cat was homozygous and affected, indicating a second cat with the disease variant from a different region of the world. The WES also supports the determination of allele frequencies of disease variants, identifying heterozygote cats for recessive diseases, such as, porphyria, Factor XII deficiency and copper metabolism ^61 71 63^. Thus, together with the WGS 99 Lives dataset, the variant frequency data can help determine the likelihood of variants being causal for diseases. The variant frequencies may also be useful for cross-species comparisons to hopefully better define variants of uncertain significance^72^.

Precision / Genomic Medicine, i.e. genomic DNA profiling, in companion animals allows veterinarians to adapt treatments to the specific animal and to the specific disease type^73^. Many rare diseases and cancers have poor prognosis, with some less than 90 days, thus, Precision / Genomic Medicine may help discover alternate and more effective treatments^74^. The Undiagnosed Diseases Program of the National Institutes of Health routinely uses WES, suggesting veterinary medicine could benefit in the same manner ^75^. WES, when compared to WGS, has proven more cost effective, more time efficient and requires fewer computing resources. Alternative uses for this exome resource could also so be developed, examples include resequencing of ancient DNA samples and important biological regions, such as the major histocompatibility complex^76^. The effective use of WES in a Precision Medicine context depends on its ability to discover disease variants. The presented feline WES capture design is robust for disease variant detection in cats. The vast majority of variants discoverable using WGS were also found using WES, known variants were identified and several novel variants were suggested and can be evaluated in detail. Importantly, based on our findings, improvements in the cat exome capture resource are also expected.

## Acknowledgments

Funding was provided in part by the Gilbreath McLorn Endowment of the MU College of Veterinary Medicine, Winn Feline Foundation / Miller Trust (MT18-009, MT19-001). We thank Thomas Juba for sample processing and consultation on variant validation. We thank Dr. Bill Murphy for advice on Y chromosome gene sequences.

## Supplementary Files

Supplementary Information: Supplementary Tables and Figures. Supplementary Data S1: Exome primary targets

Supplementary Data S2: Platform bias of genes indicated as the difference in the total number of variants in each platform. Genes are sorted by WGS – WES variants, largest to smallest.

Supplementary Data S3: The top 50 genes from Supplementary Data S2 sorted by position.

Supplementary Data S4: Sex bias of all X chromosome genes. Columns represent the mean number of SNPs per individual for a particular platform and sex. For example, “WGS.m” is the mean number of WGS variants per male individual.

Supplementary Data S5: Confirmed known variants

